# Strain variation in *Clostridioides difficile* toxin activity associated with genomic variation at both PaLoc and non-PaLoc loci

**DOI:** 10.1101/2021.12.08.471880

**Authors:** Katie Saund, Ali Pirani, D. Borden Lacy, Philip C. Hanna, Evan Snitkin

## Abstract

Clinical disease from *Clostridioides difficile* infection can be mediated by two toxins and their neighboring regulatory genes encoded within the five-gene pathogenicity locus (PaLoc). We provide several lines of evidence that the toxin activity of *C. difficile* may be modulated by genomic variants outside of the PaLoc. We used a phylogenetic tree-based approach to demonstrate discordance between toxin activity and PaLoc evolutionary history, an elastic net method to show the insufficiency of PaLoc variants alone to model toxin activity, and a convergence-based bacterial genome-wide association study (GWAS) to identify correlations between non-PaLoc loci with changes in toxin activity. Combined, these data support a model of *C. difficile* disease wherein toxin activity may be strongly affected by many non-PaLoc loci. Additionally, we characterize multiple other *in vitro* phenotypes relevant to human infections including germination and sporulation. These phenotypes vary greatly in their clonality, variability, convergence, and concordance with genomic variation. Lastly, we highlight the intersection of loci identified by GWAS for different phenotypes and clinical severity. This strategy to identify the overlapping loci can facilitate the identification of genetic variation linking phenotypic variation to clinical outcomes.

**IMPORTANCE:** *Clostridioides difficile* has two major disease mediating toxins, A and B, encoded within the pathogenicity locus (PaLoc). In this study we demonstrate via multiple approaches that genomic variants outside of the PaLoc are associated with changes in toxin activity. These genomic variants may provide new avenues of exploration in the hunt for novel disease modifying interventions. Additionally, we provide insight into the evolution of several additional phenotypes also critical to clinical infection such as sporulation, germination, and growth rate. These *in vitro* phenotypes display a range of responses to evolutionary pressures and as such vary in their appropriateness for certain bacterial genome wide association study approaches. We used a convergence-based association method to identify the genomic variants most correlated with both changes in these phenotypes and disease severity. These overlapping loci may be important to both bacterial function and human clinical disease.

## INTRODUCTION

*Clostridioides difficile* is a toxin-producing, healthcare-associated bacterial pathogen. It exhibits extensive genetic variation due to its highly mobile genome, a large pangenome, and a most recent common ancestor for clades C1-5 dating back approximately 3.89 million years (1, 2). Such genomic variability has enabled *C. difficile* adaptation to multiple host species and to spread among humans in both nosocomial and community contexts (3). Underlying this genetic variation, is phenotypic variation in many traits including toxin production, sporulation, germination, growth, and virulence (4). This genetic and phenotypic variation has led many to ask whether different genetic backgrounds of *C. difficile* may differ in their propensity to cause severe infections. To this end, many studies have sought to identify key genetic traits harbored by putative hypervirulent strains, such as Ribotype 027 (RT027). Despite this interest and intense study, the genetic basis for variation in phenotypes relevant to the *C. difficile* infection lifecycle remains limited.

Disease during *C. difficile* infection is mediated by extracellular toxins, primarily Toxins A (TcdA) and B (TcdB), which damage the cytoskeletons of intestinal cells leading to cell death and gut inflammation. These two toxins are large proteins with four domains: glucosyltransferase, autoprotease, pore-forming, and C-terminal combined repetitive oligopeptides (CROPs) (5). Toxins A and B are both located within the pathogenicity locus (PaLoc) with three other genes: *tcdR, tcdC*, and *tcdE. tcdR* is a positive regulator of *tcdA* and *tcdB* and encodes an RNA polymerase factor (6). *tcdC* may be a negative regulator of *tcdR* (6). *tcdE* encodes a holin-like protein and may contribute to toxin secretion (7). Many factors and systems are implicated in PaLoc regulation including growth phase, access to specific metabolites, sporulation, quorum sensing, and some flagellar proteins (8). In addition to toxin production, other phenotypes may influence *C. difficile* disease severity or transmission, including sporulation, germination, and growth (9–11).

Approaches for uncovering the genomic determinants of bacterial phenotypes, such as toxin activity, include *in vitro* assays, comparative genomics, and bacterial genome-wide association studies (bGWAS). An advantage of bGWAS is the ability to sift through existing genetic variation in bacterial populations to identify variants associated with natural phenotypic variation. In this way, bGWAS can provide insight into phenotypic evolution, and enable the identification of variants of interest that mediate modulation of clinically relevant phenotypes, such as virulence (12). Here, we capitalized on a diverse collection of over 100 *C. difficile* isolates for which multiple phenotypes had previously been characterized (4). We performed whole genome sequencing and used bGWAS to uncover novel genotype-phenotype associations. We explore these genotype-phenotype associations and describe the phenotype variation through phylogenetic and evolutionary analyses. Our analyses reveal the influence of genetic variation on phenotypic variation and help illuminate factors that may be influencing clinical disease.

## RESULTS

### Distinct evolutionary trajectories of clinically relevant *C. difficile* phenotypes

To improve our understanding of the evolution of phenotypic diversity in *C. difficile* we performed whole-genome sequencing on a clinical isolate collection that had previously been assayed for toxin activity, two measures of germination, two measures of sporulation, and growth rate (4, 10). Overlaying these phenotypes on a whole-genome phylogeny revealed distinct patterns for each phenotype (Fig. 1). Toxin activity and germination in Tc and Gly are clonal phenotypes that show stable inheritance within lineages, as evidenced by high phylogenetic signal (Fig. 2A). For example, toxin activity displays clonal lineages with uniformly high (e.g. RT027) and low (e.g. RT014) toxin activity (Fig. 1). In contrast, germination in Tc and growth rate are less clonal, with extensive variation even within clonal lineages (Fig. 1, 2A). Finally, the two sporulation phenotypes show the least clonality, with virtually no clustering on the phylogeny (Fig. 1,2A). Overall, the range in clonality and phylogenetic signal observed for these phenotypes suggests that despite all being central to the *C. difficile* life cycle, that they are shaped by different evolutionary pressures.

**FIG 1.**
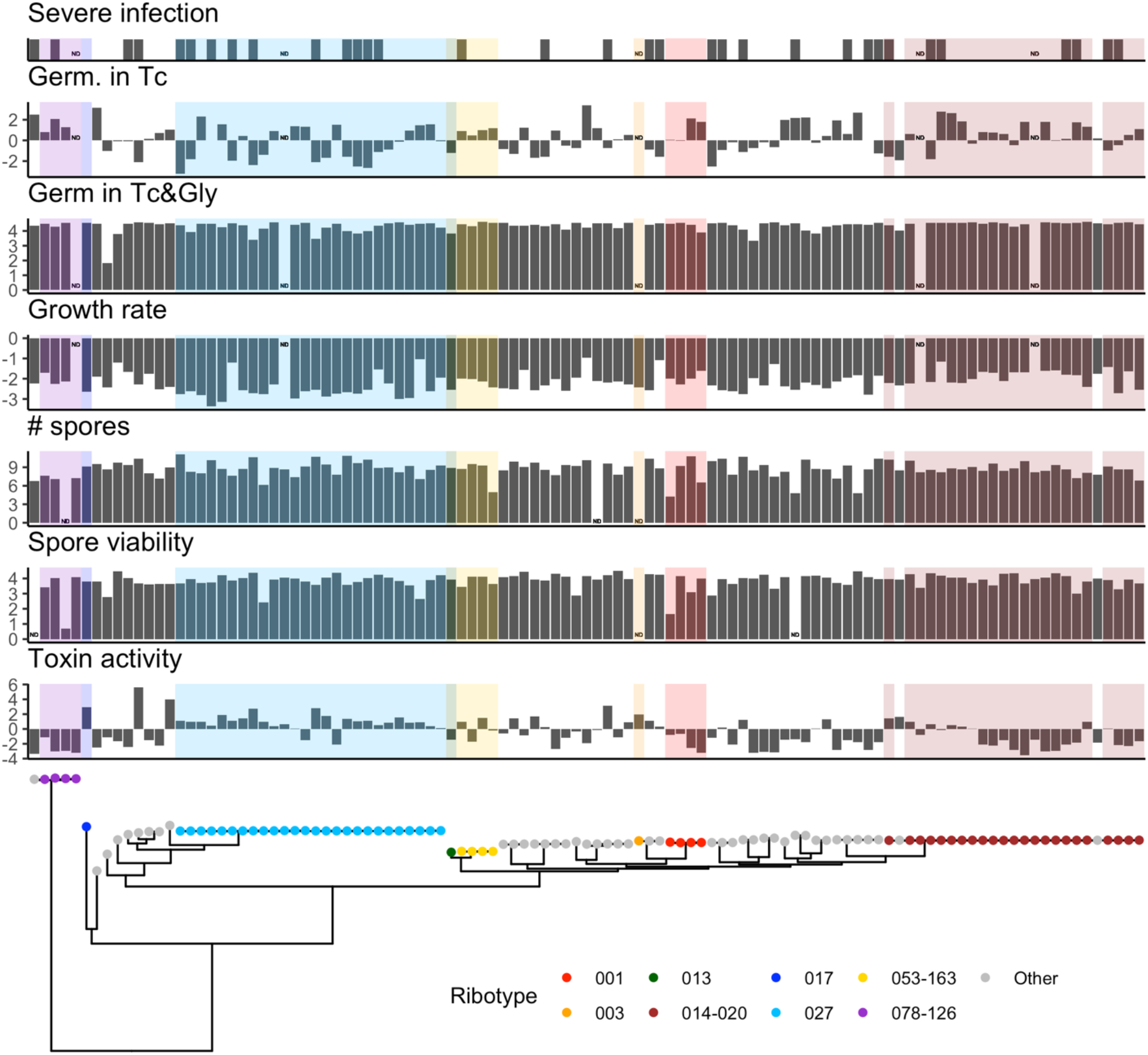
Clinical *C. difficile* sample phenotypes aligned with the phylogenetic tree. Color indicates ribotype. ND = No data. *In vitro* phenotypes were log transformed. Infections were classified as severe or not severe.

**FIG 2.**
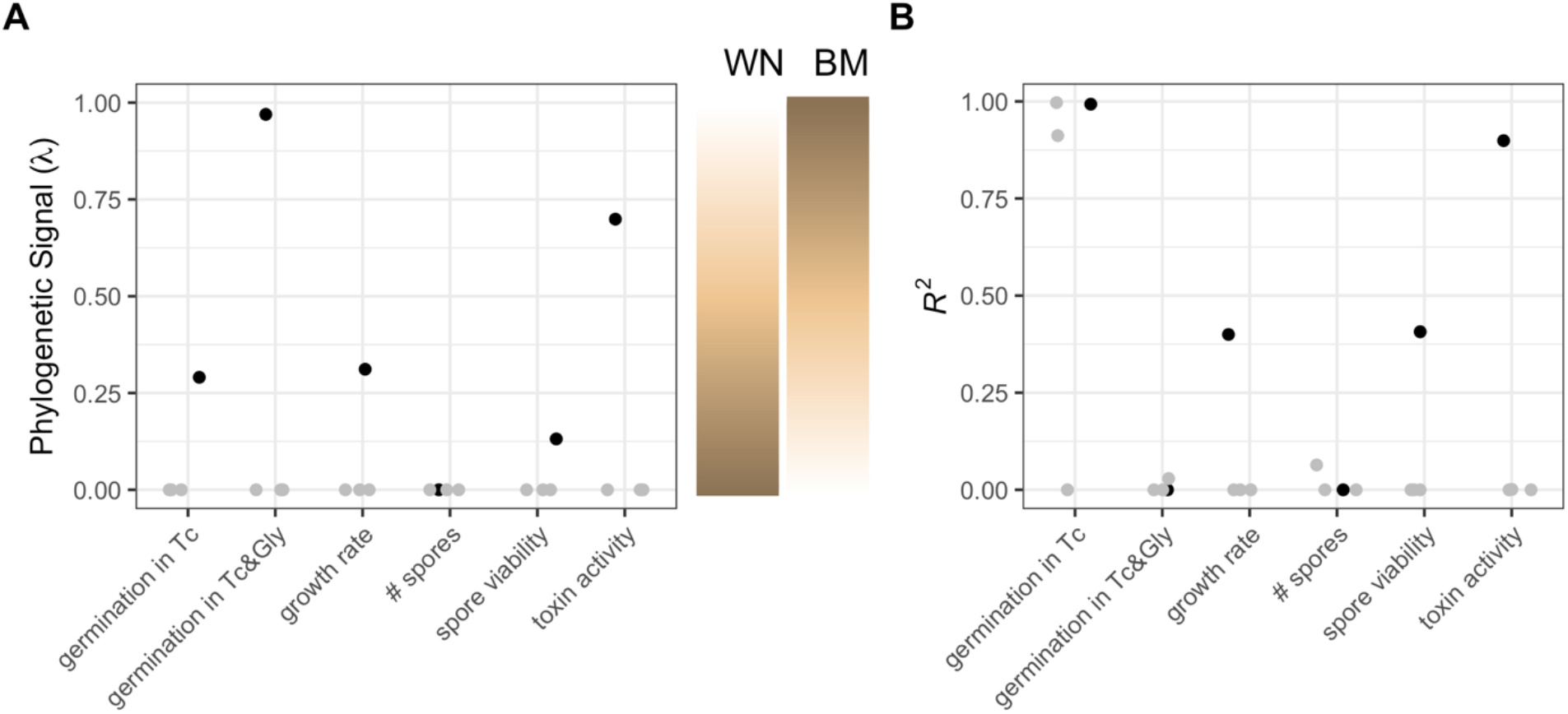
Phenotype phylogenetic signal and genomic model. (A) The phylogenetic signal of each phenotype (black) and its negative controls (grey). WN = white noise. BM = Brownian motion. (B) Elastic nets modeling each phenotype, with high R^2^ values indicating that the phenotype is strongly predicted by genetic variation in SNPs. Synonymous SNPs were excluded from this analysis.

In addition to varying in their clonality, the six phenotypes show distinct differences in their overall degree of variation (Table 1). Toxin activity had the largest dispersion with a geometric coefficient of variation of 5.4. The combination of high clonality and high dispersion in toxin activity suggests that *C. difficile* may have evolved multiple successful toxin strategies or have different evolutionary trajectories that are difficult to escape once begun. In contrast, the near uniformity observed in germination in Tc and Gly, could indicate either strong stabilizing selection or inadequate precision of the assay.

**TABLE 1.** Dispersion (geometric coefficient of variation) and convergence (ratio metric of convergence) of the log transformed phenotypes.

|  | Germination<br>in Tc | Germination<br>in Tc & Gly | Growth<br>rate | Total<br>spores | Viable<br>spores | Toxin<br>activity |
| --- | --- | --- | --- | --- | --- | --- |
| Geometric<br>coefficient<br>of variation | 2.8 | 0.4 | 0.5 | 2.2 | 0.6 | 5.4 |
| Ratio metric<br>of<br>convergence | 46.8 | 18.0 | 43.0 | 38.7 | 27.3 | 33.0 |

### Phenotypes vary with respect to their association with genetic variation

Next, we sought to understand the degree to which phenotypic variability in this dataset is genetically encoded. The phenotype best modeled by genomic variants is toxin activity with *R*^*2*^ = 0.90 (Fig. 2B). Growth rate, both sporulation phenotypes, and gemination in Tc and Gly have much lower *R*^*2*^ values, all *R*^*2*^ < 0.50. Germination in Tc has a high *R*^*2*^ value, *R*^*2*^ = 0.99, but this finding appears to be spurious as two of the three negative controls using randomly permuted data have similarly high *R*^*2*^: 0.00, 0.91, and 1.00. The germination and number of spores phenotypes are so poorly encoded by genomic variation that it is suggestive that the assays may lack sufficient precision to capture relevant strain variation, while toxin activity appears far more genetically deterministic.

### Phenotypes show a range in their level of phylogenetic convergence

A striking feature observed when overlaying the phenotype panel on the whole-genome phylogeny was variation in the frequency of convergence of high or low phenotype values. Convergence, the independent evolution of a trait, may imply the existence of environmental pressures that select towards a specific value or constrain the phenotype’s value. To quantify convergence of the different phenotypes we employed the ratio metric, where a higher ratio metric value suggests more episodes of convergence. Germination in Tc has the most convergence, 46.8. The germination in Tc and Gly and spore viability phenotypes have the least convergence, 18.0 and 27.3 respectively. These low values may be driven in part by the lack of dispersion in the phenotype values. The remaining phenotypes demonstrate intermediate levels of phylogenetic convergence. Below we seek to exploit the high level of convergence in certain phenotypes to identify genetic drivers of their variation.

### Identifying genetic variation associated with phenotypic variation through genome-wide association study

Having observed differences in the evolutionary patterns of different phenotypes, we next sought to identify the specific genetic variation that may be underlying phenotypic variation by performing a genome-wide association study (GWAS) for each phenotype. Due to the high convergence in several of the phenotypes (Table 1) and extensive genetic variation in our isolate collection, we opted for a convergence-based GWAS approach that could identify variants of interest by their non-random co-convergence with a phenotype. The genotypes tested included approximately 69,600 SNPs, 8,400 indels, and 7,500 accessory genes. Significantly associated variants were identified for growth rate, number of spores, toxin activity, germination in Tc, and severity.

#### Overlapping GWAS results

Despite the phenotypes showing distinct evolutionary patterns, we first explored whether there was evidence of overlap in the genetic circuits modulating the different traits. We cataloged the extent of this overlap by counting the number of intersecting genomic loci with both high significance and convergence in each pair of GWAS results. Three of the four phenotypes shared more hits with the severe infection GWAS results than expected by chance via a permutation test (Fig. 3A). Toxin activity and severe infection have the most overlap with 7 shared loci. These shared loci include six accessory genes and a frameshift mutation at Glycine 209 in flagellar hook-associated protein 2 (*fliD)* (Fig. 3B). The *fliD* finding is consistent with known co-regulation that occurs between flagellar and toxin systems in *C. difficile* that is mediated in part by SigD, a sigma factor that binds to a *tcdR* promoter region and positively regulates *tcdR* (13).

**FIG 3.**
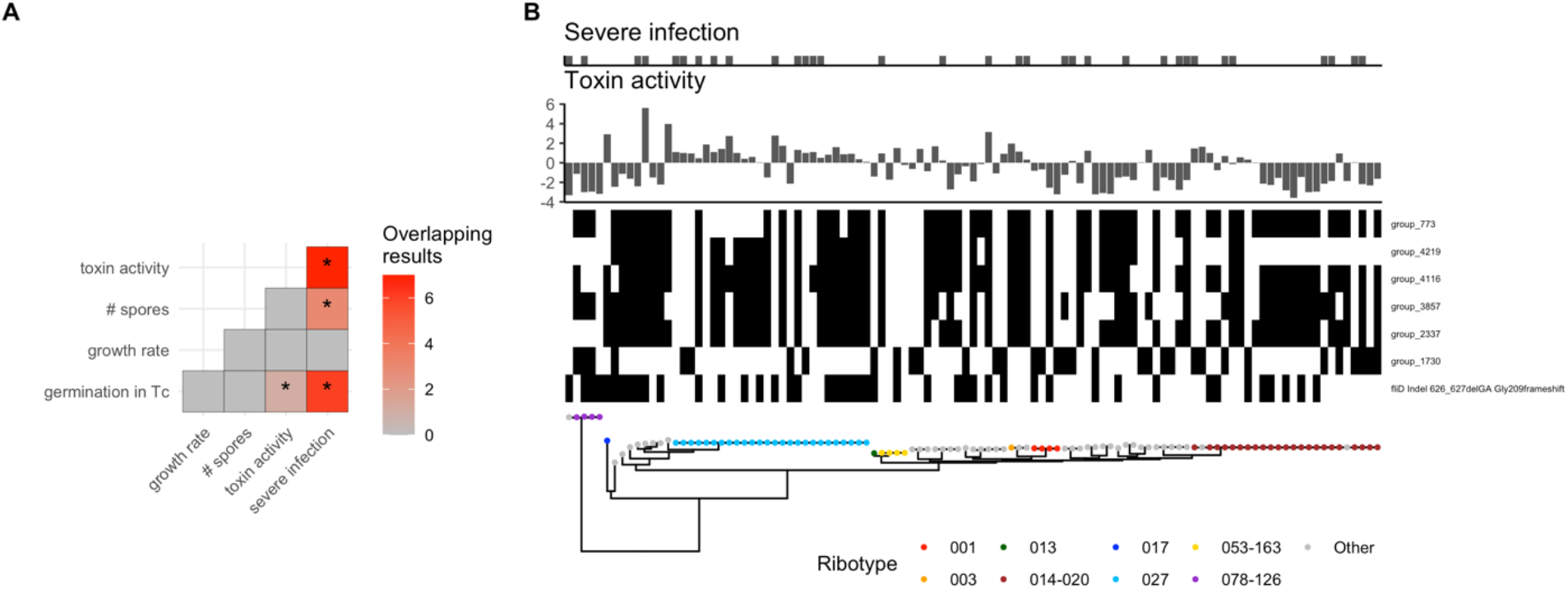
Overlapping GWAS results. (A) Heatmap indicates the number of shared GWAS results with significant *P*-values and high levels of convergence in the Continuous Test (continuous phenotypes) or Synchronous Test (Severity). Asterisks indicate significantly more overlapping results than expected by chance (*P* < 0.05). The two phenotypes lacking any GWAS results with significant *P*-values and high levels of convergence were excluded. (B) Shared hits between the toxin activity and severe infection GWAS. Top: phenotype. Center: heatmap indicating the presence of loci with both significant *P*-values and high levels of convergence in both toxin activity and severity GWAS results. Bottom: phylogenetic tree labeled by ribotype.

### Genetic variation associated with modulation of toxin activity

For the remainder of our analysis, we focused on understanding genetic variation associated with variation in toxin activity. In addition to the central role of toxin in *C. difficile* disease, our decision to focus on toxin was motivated by it being the phenotype being best explained by genetic variation in sequenced strains (Figure 2B). In the following sections we examine variants playing a key role in modulating toxin activity.

The toxin activity GWAS identified many genomic variants both significantly associated with toxin activity changes and had high levels of convergence (Fig. 4A). As the PaLoc encodes toxin genes and regulators we expected that variants located within the PaLoc would be significantly associated with toxin activity and used this as a positive control for our analysis. Consistent with this, we observed PaLoc variants in the pool of significant results associated with toxin activity. Eighty-seven of the 220 loci significantly associated with toxin activity occur in the PaLoc. Given that the toxin activity assay used is a measure of Toxin B activity it is particularly reassuring that these 87 PaLoc loci include 75 *tcdB* variants and 2 *tcdR*-*tcdB* intergenic region variants (Fig. 4B). Indeed, these variants are a significant enrichment compared to the number of variants within or flanking *tcdB* that are expected by chance using a permutation approach, *P* = 0.0001 (median = 1; range = 0-10). *tcdB* variants were found in all four protein domains, but the significantly associated variants are mostly found within the glucosyltransferase and autoprotease domains (Fig. 4B). Certain significant missense variants within *tcdB* have plausible functional impacts on Toxin B such as an adenosine to cytosine transversion at position 1967 which changes an aspartic acid to alanine; this mutation occurs near the zinc binding site and could theoretically affect toxin autoprocessing within the host cell. Of the 15 tested variants that occur within the *tcdR*-*tcdB* intergenic region, 5 were significantly associated with toxin activity. Three of these variants occur within a *tcdB* promoter suggesting a potential role in modulating sigma factor binding and therefore altering *tcdB* transcription. A notable lack of association was observed between an adenosine deletion at nucleotide 117 in *tcdC* that has been suggested to cause increased toxin production in RT027 (14). This deletion was found in all 26 RT027 samples as well as in 3 additional samples (“Other” ribotype) but did not reach significance in the GWAS, *P*=0.95.

**FIG 4.**
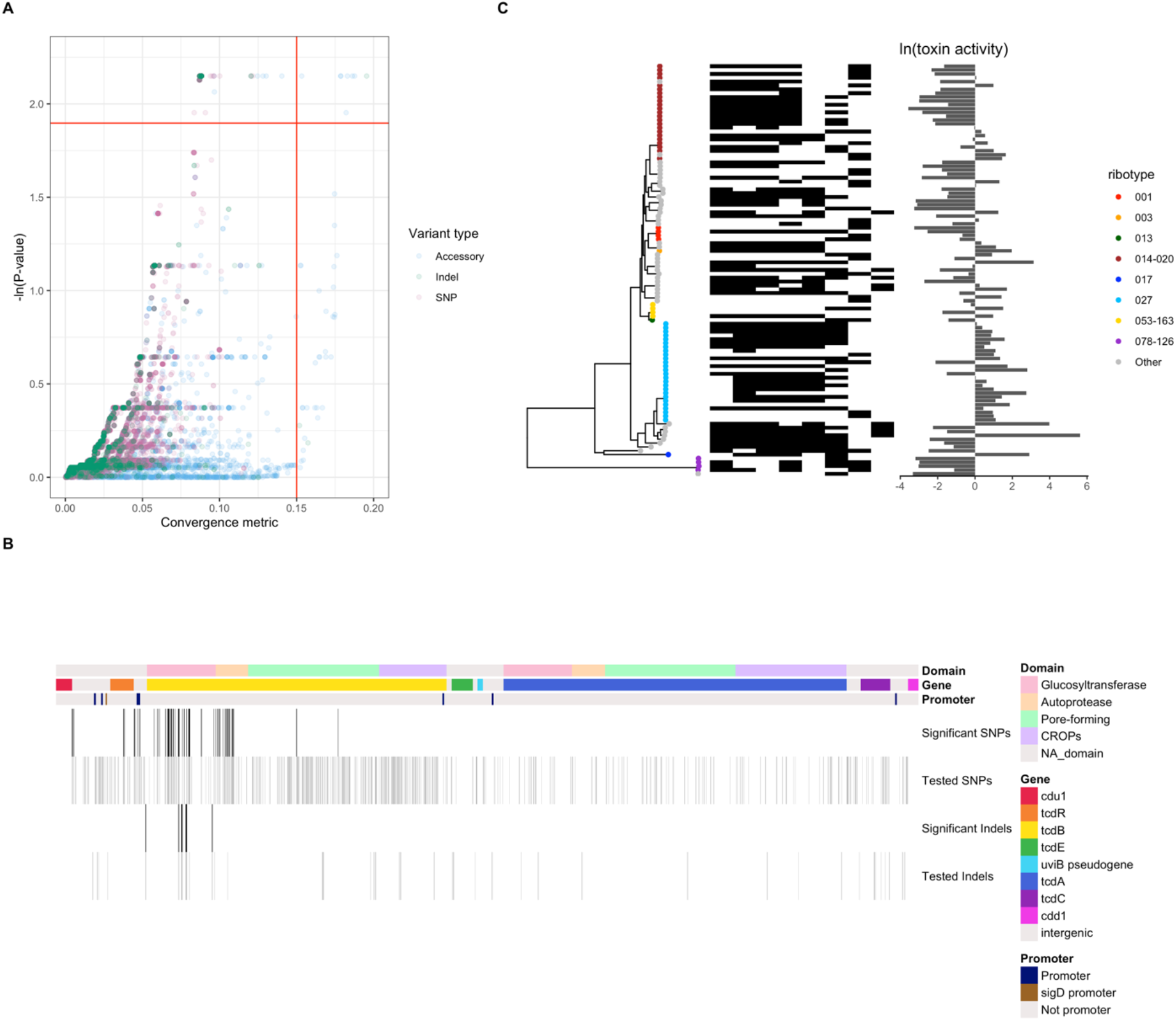
Genome-wide association study identified genomic variants associated with toxin activity variation. (A) GWAS results. Tested loci are either accessory genes (blue; N=4,352), SNPs (pink; N=12,167), or indels (green; N=1,843). The red horizontal line indicates a False Discovery Rate of 15%. The red vertical line separates low vs high convergence. (B) Significantly associated loci from GWAS located in the PaLoc. Of the 633 PaLoc variants (SNP N=563, Indel N=70) tested by GWAS only the variants significantly associated are plotted as vertical bars (SNP N=71, Indel N=16). Top annotation: toxin protein domains in *tcdB*. Center annotation: gene. Bottom annotation: promoter locations. (C) Left: phylogenetic tree labeled by ribotype. Center: heatmap indicating the presence of loci significantly associated with toxin activity and with high convergence. Right: toxin activity.

Next, we sought to generate hypotheses about new associations between genomic variants and toxin activity that reside outside of the PaLoc. The variants that were significant and had high ε, a metric of shared genotype-phenotype convergence, are cataloged in File S1 and plotted in Fig. 4C (15). A single ε value captures the number of tree edges where both a genotype is mutated and the toxin activity value has a large change. ε values close to zero suggest that the genotype mutates on very few edges where the toxin activity changes drastically. The loci associated with changes in toxin activity are present in multiple, independent lineages (Fig. 4C). The previously mentioned frameshift mutation in *fliD* is the variant most strongly associated with changes in toxin activity when ranked by ε then *P*-value. The next most strongly associated variant maps to CD630_02364 which is annotated as a putative signaling protein in the reference genome CD630. The other most strongly associated variants are poorly annotated accessory genes. The significant accessory genes identified by this analysis may yield profitable results in mechanistic studies dissecting *C. difficile* toxin activity and could be prioritized for further characterization.

### Genetic variation at PaLoc accounts for only half of phenotypic variation in toxin activity

The GWAS identified both PaLoc and non-PaLoc loci correlated with variation in toxin activity. To understand the relative contribution of genetic variation in the PaLoc to variation in toxin activity we employed an elastic net approach. Models of toxin activity constructed with different subsets of variants found that PaLoc variants and *tcdB* variants have similar abilities to model toxin activity, *R*^*2*^ = 0.48 and *R*^*2*^ = 0.46 respectively (Fig. 5A). However, variants from the whole genome build a more accurate model of toxin activity, *R*^*2*^ = 0.90 (Fig. 5A). Of the 634 variants in the PaLoc, 404 (64%) occur in *tcdB* or in its flanking intergenic regions; in the best performing elastic net model derived from PaLoc variants, 34/61 (56%) of the variants are mutations in *tcdB* or its flanking regions. In the whole genome model only 17/1795 (1%) of variants occur in *tcdB* or its flanking regions.

**FIG 5.**
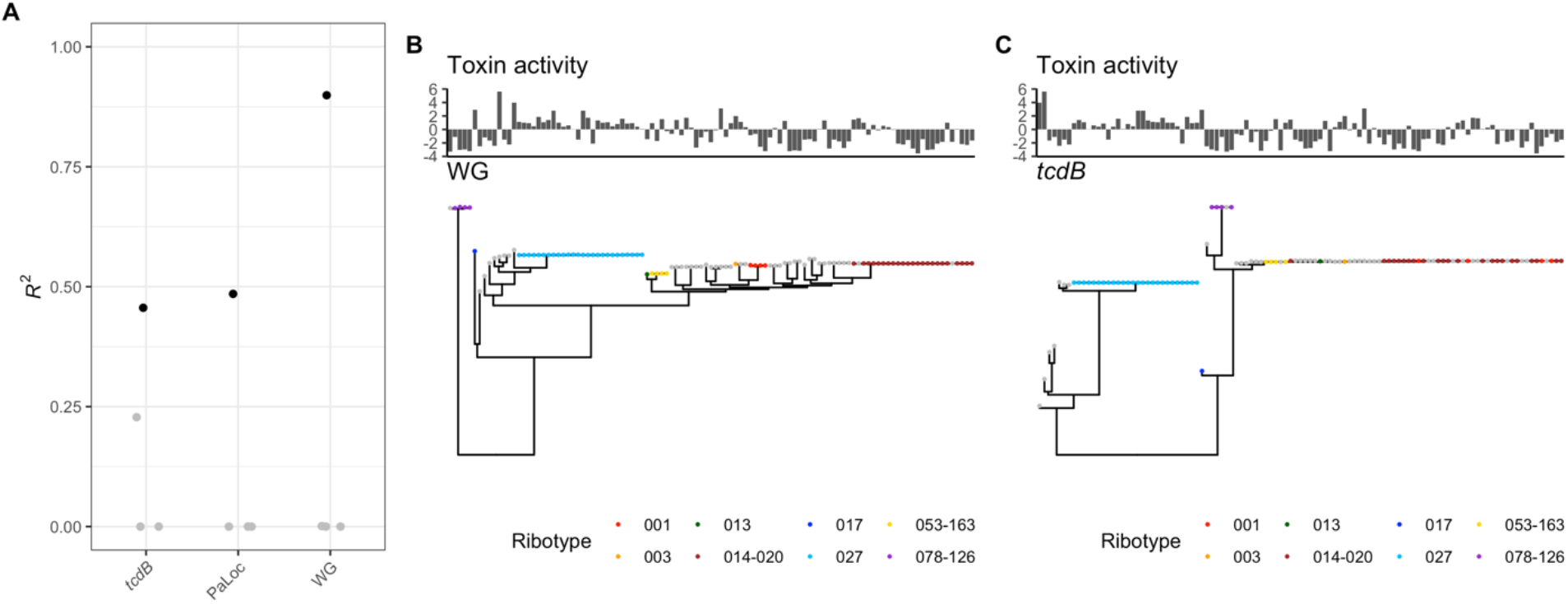
*tcdB* variation does not fully model toxin activity. (A) Elastic net model performance of toxin activity. Models were built from *tcdB* variants, PaLoc variants, or whole genome (WG) variants. Toxin activity with a tree built from (B) the whole genome or (C) just *tcdB*.

To assess the predictive capacity of PaLoc variation in a different way, we compared phylogenetic trees built from whole genome variants and the *tcdB* gene. As there is far less variation in *tcdB* than the whole genome, we observe many polytomies in the *tcdB* gene tree and none in the whole genome tree (Fig 5B,C). While the *tcdB* gene tree’s toxin activity is better modeled by Brownian motion, λ = 0.94, than in the whole genome tree, λ = 0.75 (Fig. 2A), there remains much toxin activity variation unexplained by tree structure. Given the unexplained toxin activity variation on the *tcdB* gene tree and variation not captured in the toxin activity elastic net model, we conclude that while *tcdB* gene variation is likely an important mediator in the evolution of toxin activity, other loci play a key role as well. Finally, the whole genome model suggests that many loci besides *tcdB* may affect *C. difficile* toxicity and therefore a wider lens for examining genetic influences on toxicity will be fruitful.

## DISCUSSION

*C. difficile* is a genetically diverse pathogen, with extensive variation in both its core and accessory genome. Currently, we have a limited understanding of the functional impact of most of this variation and how it relates to *C. difficile* infection. Here, we attempted to improve our understanding of the genotype to phenotype map in *C. difficile* by analyzing variation in clinically relevant phenotypes in the context of *C. difficile* genomic variants. We observe that despite their central role to the *C. difficile* transmission and infection cycle sporulation, germination, growth and toxin activity show distinct evolutionary trajectories. Focusing on the phenotype thought to be most closely linked to virulence, we observe that toxin activity is highly clonal, with lineages tending to either possess high or low toxin activity. Consistent with prior reports we find that variation in toxin activity can be modulated by variants in the PaLoc, however we find that more than 50% of phenotypic variation is associated with genetic variation outside of the PaLoc.

Our exploration of these *C. difficile* phenotypes revealed a broad range of clonality, dispersion, association with genomic variation, and convergence. As such, each phenotype appears to be shaped by different selection forces. The existence of phenotypes that show no association with the recombination filtered phylogeny could indicate either a lack of precision in the laboratory assay or a strong role for recombinant genomic regions in shaping these phenotypes. We focused our analysis on toxin activity, in part, because of the precision of the *in vitro* assay results and its high degree of genetic determinism. Regardless of the basis for the lack of phylogenetic signal in some of the non-toxin phenotypes, these results show how overlaying phenotypic variation on whole-genome phylogenies provides useful context for interpreting and scrutinizing experimental measurements, and in this case clearly demonstrates the rich and varied patterns of evolution among *C. difficile* strains.

Toxigenic bacterial species that require live transmission may undergo strong selective pressure to promote host survival and therefore bias towards lower toxin activity (16). In contrast, sporogenic *C. difficile* can survive and transmit even after the host dies; this may reduce the strength of selection on toxin activity and therefore many different toxin strategies are successful. Indeed, there are prolific toxigenic and non-toxigenic strains of *C. difficile*. Additionally, the species has had multiple independent losses of the PaLoc (17), with our results indicating that even strains harboring an intact PaLoc may evolve to have decreased toxin activity. The *C. difficile* strains with high toxin activity may have success by shaping a hostile metabolic state in the host gut that these bacteria are able to uniquely exploit (18) or its more severe, inflammatory infection which results in diarrhea and therefore increased transmission. This then raises the question of what the selective pressure for lower toxin activity may be. One possibility is that toxin activity itself may not be the most critical aspect of the toxin upon which evolution is acting, with other aspects such as toxin immunogenicity potentially evoking a stronger selection pressure. Toxin that evades immune recognition could lead to longer infections and therefore increased transmission, so the strongest selective pressure may be at the surface domains of the toxin proteins rather than on regulators of toxin activity (19). For example, we observed multiple missense variants on the surface of tcdB in this isolate collection, including a glutamic acid 329 to glycine missense variant and threonine 430 to alanine.

Our study has several important limitations. First, the limited sample size of this *C. difficile* collection could lead to underreporting of clonality of some phenotypes for underrepresented ribotypes and limits power to detect variation with smaller phenotypic impacts. Second, many genomic features such as copy number variants, large structural variants, and plasmids were not included in our GWAS or elastic net models, therefore these analyses are missing some genome encoded information. Similarly, we did not consider the complexity of epistatic interactions between genomic variants on phenotypes.

A replication study in a second *C. difficile* cohort in which the toxin activity assay and GWAS is repeated could help prioritize the genomic variants more likely to be causal of changes in toxin activity. The loci identified in both this study and the proposed study would be higher confidence candidates for experiments that examine the effect of those potential variants on toxin activity. Additional studies investigating *C. difficile in vitro* phenotypes from an evolutionary perspective would help to prioritize the phenotypes that may offer the most insight into the success and regulation of certain strains.

## MATERIAL AND METHODS

### Study population and *in vitro* characterization

The University of Michigan Institutional Review Board approved all sample and clinical data collection protocols used in this study (HUM00034766). Where applicable, written, informed consent was received from all patients prior to inclusion in this study. Stool samples were collected from a cohort of 106 Michigan Medicine patients with *C. difficile* infection from 2010-2011, which included all severe cases during the study period (4, 10). Cases were classified as severe if the infection required ICU admission or interventional surgery, or if the patient died within 30 days of infection diagnosis. A clonal spore stock from each patient was used for ribotyping and *in vitro* studies. Previous experiments characterized the germination in taurocholate (TC; %), and germination in Tc and glycine (Gly; %), maximum growth rates (OD_600_/hour), total spore production (heat resistant colon forming units per ml), viable spores (%), and equivalent toxin B activity (ng/ml) (4, 10). Taurocholate is a physiologic bile salt known to cause *C. difficile* germination; glycine can increase germination with taurocholate (20). Samples were classified as severe infections if they were collected from a patient whose *C. difficile* infection required ICU admission or interventional surgery, or if the patient died within 30 days of infection diagnosis (4, 10).

### Genomic analysis

The spore stocks were grown in an anaerobic chamber overnight on taurocholate-coition-cycloserine-fructose agar plates. The next day a single colony of each sample was picked and grown in Brain Heart Infusion medium with yeast extract liquid culture media overnight. The vegetative *C. difficile* cells were pelleted by centrifugation, washed, and then total genomic DNA was extracted. Genomic DNA extracted with the MoBio PowerMag Microbial DNA Isolation Kit (Qiagen) from *C. difficile* isolates (N=108) was prepared for sequencing using the Illumina Nextera DNA Flex Library Preparation Kit. Sequencing was performed on either an Illumina HiSeq 4000 System at the University of Michigan Advanced Genomics Core or an Illumina MiSeq System at the University of Michigan Microbial Systems Molecular Biology Laboratories. Quality of reads was assessed with FastQC v0.11.9 (21). Adapter sequences and low-quality bases were removed with Trimmomatic v0.36 (22). Variants were identified by mapping filtered reads to the CD630 reference genome (GenBank accession number AM180355.1) using bwa v0.7.17 (23), removing polymerase chain reaction duplicates with Picard 2.21.7 (24), removing clipped alignments using Samclip 0.4.0 (25), and calling variants with SAMtools v1.11and bcftools (26). Variants were filtered from raw results using GATK’s VariantFiltration v3.8 (QUAL, >100; MQ, >50; >=10 reads supporting variant; and FQ, <0.025) (27). SNPs and indels were referenced to the ancestral allele using snitkitr v0.0.0.9000 (28). Pangenome analysis was performed with roary (29). Accessory genes annotations were assigned by prokka v1.14.5 (30).

### Data availability

Sequence data are available under Bioproject PRJNA594943. Details on sequenced strains are available in File S2. Sequences for genes identified by roary are available in File S3.

### Phylogenetic analysis

Consensus files generated during variant calling were recombination filtered using Gubbins v3.0.0 (31). The alleles at each position that passed filtering were concatenated to generate a non-core variant alignment relative to the CD630 reference genome. Alleles that did not pass filtering were considered unknown (denoted as N in the alignment). Variant positions in the alignment were used to reconstruct a maximum likelihood phylogeny with IQ-TREE v1.5.5 using ultrafast bootstrap with 1,000 replicates (32, 33). ModelFinder limited to ascertainment bias-corrected models was used to identify the best nucleotide substitution model (34). The tree was midpoint rooted. The *tcdB* multiple sequence alignment was built by PRANK v.170427 using only the *tcdB* gene and the resulting tree was midpoint rooted (35). The trees are available in Files S4 and S5.

### Genome-wide association studies

GWAS were performed with hogwash v1.2.4 (15). Phenotype data were natural log transformed. Hogwash settings: bootstrap threshold=0.95, permutations=10,000, false discovery rate=15%. The analysis included SNPs, indels, and accessory genes. The intersection of hogwash results was restricted to results with ε > 0.15 and *P*-value < 0.15. Only SNPs classified as having “Moderate”, “High”, or “Modifier” impact by SnpEff v4.3.1 were included (36).

### Data analysis

Data analysis with R v3.6.2 (37) was performed with following packages: ape v5.3 (38), aplot v0.0.6 (39), data.table v1.12.8 (40), ggtree v2.0.4 (41), ggpubr v0.4.0 (42), pheatmap v1.0.12 (43), phytools v0.6-99 (44), tidyverse v1.3.0 (45). Conda v4.9.2 was used to maintain working environments (46). Analysis code is available at https://github.com/katiesaund/cdifficile_gwas.

### Permutation testing

The empirical *P*-value for enrichment of toxin variants in the significant GWAS results and shared results in the overlapping GWAS section were generated via permutation testing. This approach generates a *P*-value by comparing the observed number of events in the data to a distribution of the number of events simulated under the null hypothesis. The null distribution was generated from random sampling without replacement repeated in 10,000 trials (toxin variants) or 1,000 trials (overlapping hits). Multiple testing correction was applied to the overlapping hits analysis using Bonferroni correction.

### Convergence analysis

We calculated the degree of convergence of each phenotype using the ratio method (47), which is the ratio of two sample’s pairwise patristic distance divided by their pairwise phenotypic distance. We report the average of the scaled pairwise branch length distance (patristic distance) divided by scaled pairwise phenotypic distance for each phenotype. A high value suggests an episode of convergence.

### Geometric coefficient of variance

We calculated the dispersion of each phenotype as defined by the geometric coefficient of variance: 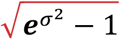 where α is the standard deviation of the log transformed data.

### Phylogenetic signal

Phylogenetic signal is a metric that captures the tendency for closely related samples on a tree to be more similar to each other than they are to random samples on the tree. We calculated phylogenetic signal for each continuous phenotype using Pagel’s λ (48). Note that a phenotype that is modeled well by Brownian motion has a λ near 1 while a white noise phenotype has a λ near 0 (48). Negative controls for the phenotypes were created by randomly redistributing each phenotype on the tree.

### Elastic net modeling

We calculated the degree of genetic encoding of each phenotype by modeling a phenotype from genomic variants using elastic net regularization as implemented by pyseer. Pyseer v1.3 command line arguments: --wg enet --n-fold 10 (49). SNPs, indels and accessory genes were all used to model all continuous phenotypes. For all elastic net models only SNPs classified as having “Moderate”, “High”, or “Modifier” impact by SnpEff were included (36). Toxin activity was additionally modeled by 1) a model built from just PaLoc SNPs and indels and 2) a model built from just *tcdB* SNPs and indels. To determine the value of α, a parameter which controls the ratio of L1 and L2 regularization in the model, five α values were tested for each model: 0.01, 0.245, 0.500, 0.745, and 0.990. The model results with the highest *R*^*2*^ value were reported. The best α for models of germination in Tc, germination in Tc and Gly, total spores, and toxin activity (all variants) is 0.01. The best α for models of viable spores and growth rate is 0.245. The best α to model toxin activity (*tcdB*) is 0.500. The best α to model toxin activity (PaLoc) is 0.745. Negative controls for the phenotypes were created by randomly redistributing each phenotype on the tree.

## SUPPLEMENTAL MATERIAL

File S1: Toxin GWAS results. A table with variant name, p-value, and epsilon value. File S2: Bioproject details for the sequenced strains.

File S3: Sequences of the genes identified by roary. File S4: Phylogenetic tree.

File S5: *tcdB* gene phylogenetic tree.

## ACKNOWLEDGEMENTS

K. S. was supported by the National Institutes of Health (T32GM007544). E. S. S. and A. P. were supported by the National Institutes of Health (1U01Al124255). Work in the Lacy lab is supported by NIH AI095755 and VA BX002943.

## Notes

### Competing Interest Statement

The authors have declared no competing interest.

### Summary of Updates

Added supplementary files.

## REFERENCES

1. Mullany P, Allan E, Roberts AP. 2015. Mobile genetic elements in Clostridium difficile and their role in genome function. Res Microbiol 166:361–367.

2. Knight DR, Imwattana K, Kullin B, Guerrero-Araya E, Paredes-Sabja D, Didelot X, Dingle KE, Eyre DW, Rodríguez C, Riley T V. 2021. Major genetic discontinuity and novel toxigenic species in Clostridioides difficile taxonomy. Elife 10:1–25.

3. Knight DR, Elliott B, Chang BJ, Perkins TT, Riley T V. 2015. Diversity and evolution in the genome of Clostridium difficile. Clin Microbiol Rev 28:721–741.

4. Carlson PE, Walk ST, Bourgis AET, Liu MW, Kopliku F, Lo E, Young VB, Aronoff DM, Hanna PC. 2013. The relationship between phenotype, ribotype, and clinical disease in human Clostridium difficile isolates. Anaerobe 24:109–116.

5. Pruitt RN, Lacy DB. 2012. Toward a structural understanding of Clostridium difficile toxins A and B. Front Cell Infect Microbiol 2:28.

6. Monot M, Eckert C, Lemire A, Hamiot A, Dubois T, Tessier C, Dumoulard B, Hamel B, Petit A, Lalande V, Ma L, Bouchier C, Barbut F, Dupuy B. 2015. Clostridium difficile: New Insights into the Evolution of the Pathogenicity Locus. Sci Rep 5.

7. Govind R, Dupuy B. 2012. Secretion of Clostridium difficile Toxins A and B Requires the Holin-like Protein TcdE. PLoS Pathog 8:e1002727.

8. Martin-Verstraete I, Peltier J, Dupuy B. 2016. The regulatory networks that control Clostridium difficile toxin synthesis. Toxins (Basel). Multidisciplinary Digital Publishing Institute.

9. Burns DA, Minton NP. 2011. Sporulation studies in Clostridium difficile. J Microbiol Methods 87:133–138.

10. Carlson PE, Kaiser AM, Mccolm SA, Bauer JM, Young VB, Aronoff DM, Hanna PC. 2015. Variation in germination of Clostridium difficile clinical isolates correlates to disease severity. Anaerobe 33:64–70.

11. Tschudin-Sutter S, Braissant O, Erb S, Stranden A, Bonkat G, Frei R, Widmer AF. 2016. Growth Patterns of Clostridium difficile - Correlations with Strains, Binary Toxin and Disease Severity: A Prospective Cohort Study. PLoS One 11:e0161711.

12. Laabei M, Recker M, Rudkin JK, Aldeljawi M, Gulay Z, Sloan TJ, Williams P, Endres JL, Bayles KW, Fey PD, Yajjala VK, Widhelm T, Hawkins E, Lewis K, Parfett S, Scowen L, Peacock SJ, Holden M, Wilson D, Read TD, Van Den Elsen J, Priest NK, Feil EJ, Hurst LD, Josefsson E, Massey RC. 2014. Predicting the virulence of MRSA from its genome sequence. Genome Res 24:839–849.

13. El Meouche I, Peltier J, Monot M, Soutourina O, Pestel-Caron M, Dupuy B, Pons JL. 2013. Characterization of the SigD regulon of C. difficile and its positive control of toxin production through the regulation of tcdR. PLoS One 8:1–17.

14. Warny M, Pepin J, Fang A, Killgore G, Thompson A, Brazier J, Frost E, McDonald LC. 2005. Toxin production by an emerging strain of Clostridium difficile associated with outbreaks of severe disease in North America and Europe. Lancet 366:1079–1084.

15. Saund K, Snitkin ES. 2020. Hogwash: three methods for genome-wide association studies in bacteria. Microb Genomics https://doi.org/10.1099/mgen.0.000469.

16. Rudkin JK, McLoughlin RM, Preston A, Massey RC. 2017. Bacterial toxins: Offensive, defensive, or something else altogether? PLoS Pathog 13:1–12.

17. Mansfield MJ, Tremblay BJM, Zeng J, Wei X, Hodgins H, Worley J, Bry L, Dong M, Doxey AC. 2020. Phylogenomics of 8,839 Clostridioides difficile genomes reveals recombination-driven evolution and diversification of toxin A and B. PLoS Pathog 16:1–24.

18. Fletcher JR, Pike CM, Parsons RJ, Rivera AJ, Foley MH, McLaren MR, Montgomery SA, Theriot CM. 2021. Clostridioides difficile exploits toxin-mediated inflammation to alter the host nutritional landscape and exclude competitors from the gut microbiota. Nat Commun 12:1–14.

19. Mansfield MJ, Tremblay BJ-M, Zeng J, Wei X, Hodgins H, Worley J, Bry L, Dong M, Doxey AC. 2020. Phylogenomics of 8,839 Clostridioides difficile genomes reveals recombination-driven evolution and diversification of toxin A and B. Biorvix.

20. Sorg JA, Sonenshein AL. 2008. Bile salts and glycine as cogerminants for Clostridium difficile spores. J Bacteriol 190:2505–2512.

21. Andrews Si. 2010. FastQC: a quality control tool for high throughput sequence data.

22. Bolger AM, Lohse M, Usadel B. 2014. Trimmomatic: A flexible trimmer for Illumina sequence data. Bioinformatics 30:2114–2120.

23. Li H, Durbin R. 2009. Fast and accurate short read alignment with Burrows-Wheeler transform. Bioinformatics 25:1754–1760.

24. Broad Institute. Picard Tools.

25. Seemann T. samclip.

26. Li H, Handsaker B, Wysoker A, Fennell T, Ruan J, Homer N, Marth G, Abecasis G, Durbin R. 2009. The Sequence Alignment/Map format and SAMtools. Bioinformatics 25:2078–2079.

27. McKenna A, Hanna M, Banks E, Sivachenko A, Cibulskis K, Kernytsky A, Garimella K, Altshuler D, Gabriel S, Daly M, DePristo MA. 2010. The Genome Analysis Toolkit: A MapReduce framework for analyzing next-generation DNA sequencing data. Genome Res 20:1297–1303.

28. Saund K, Lapp Z, Thiede SN. snitkitr.

29. Page AJ, Cummins CA, Hunt M, Wong VK, Reuter S, Holden MTG, Fookes M, Falush D, Keane JA, Parkhill J. 2015. Roary: Rapid large-scale prokaryote pan genome analysis. Bioinformatics 31:3691–3693.

30. Seemann T. 2014. Prokka: Rapid prokaryotic genome annotation. Bioinformatics 30:2068–2069.

31. Croucher NJ, Page AJ, Connor TR, Delaney AJ, Keane JA, Bentley SD, Parkhill J, Harris SR. 2015. Rapid phylogenetic analysis of large samples of recombinant bacterial whole genome sequences using Gubbins. Nucleic Acids Res 43:e15.

32. Nguyen LT, Schmidt HA, Von Haeseler A, Minh BQ. 2015. IQ-TREE: A fast and effective stochastic algorithm for estimating maximum-likelihood phylogenies. Mol Biol Evol 32:268–274.

33. Thi Hoang D, Chernomor O, von Haeseler A, Quang Minh B, Sy Vinh L, Rosenberg MS. 2017. UFBoot2: Improving the Ultrafast Bootstrap Approximation. Mol Biol Evol 35:518–522.

34. Kalyaanamoorthy S, Minh BQ, Wong TKF, Von Haeseler A, Jermiin LS. 2017. ModelFinder: Fast model selection for accurate phylogenetic estimates. Nat Methods 14:587–589.

35. Löytynoja A. 2014. Phylogeny-aware alignment with PRANK. Methods Mol Biol 1079:155–170.

36. Cingolani P, Platts A, Wang LL, Coon M, Nguyen T, Wang L, Land SJ, Lu X, Ruden DM. 2012. A program for annotating and predicting the effects of single nucleotide polymorphisms, SnpEff: SNPs in the genome of Drosophila melanogaster strain w1118; iso-2; iso-3. Fly (Austin) 6:80–92.

37. R Core Team. 2018. R: A language and environment for statistical computing. 3.5.0. R Foundation for Statistical Computing, Vienna, Austria.

38. Paradis E, Schliep K. 2019. Ape 5.0: An environment for modern phylogenetics and evolutionary analyses in R. Bioinformatics 35:526–528.

39. Yu G. 2020. aplot: Decorate a “ggplot” with Associated Information. 0.0.6.

40. Dowle M, Srinivasan A. 2020. data.table: Extension of ‘data.frame’. 1.12.8.

41. Yu G, Smith DK, Zhu H, Guan Y, Lam TTY. 2017. Ggtree: an R Package for Visualization and Annotation of Phylogenetic Trees With Their Covariates and Other Associated Data. Methods Ecol Evol 8:28–36.

42. Kassambara A. 2020. ggpubr: “ggplot2” Based Publication Ready Plots. 0.4.0.

43. Kolde R. 2019. pheatmap: Pretty Heatmaps. 1.0.12.

44. Revell LJ. 2012. phytools: An R package for phylogenetic comparative biology (and other things). Methods Ecol Evol 3:217–223.

45. Wickham H. 2017. tidyverse: Easily Install and Load the “Tidyverse.” R package version 1.2.1.

46. 2016. Anaconda Software Distribution. 2-2.4.0.

47. Stayton CT. 2008. Is convergence surprising? An examination of the frequency of convergence in simulated datasets. J Theor Biol 252:1–14.

48. Pagel M. 1997. Inferring evolutionary processes from phylogenies. Zool Scr 26:331–348.

49. Lees JA, Galardini M, Bentley SD, Weiser JN, Corander J. 2018. pyseer: a comprehensive tool for microbial pangenome-wide association studies. Bioinformatics https://doi.org/10.1093/bioinformatics/bty539.

